# Combined Nanopore and Single-Molecule Real-Time Sequencing Survey of Human Betaherpesvirus 5 Transcriptome

**DOI:** 10.1101/2021.03.30.437686

**Authors:** Balázs Kakuk, Dóra Tombácz, Zsolt Balázs, Norbert Moldován, Zsolt Csabai, Gábor Torma, Klára Megyeri, Michael Snyder, Zsolt Boldogkői

## Abstract

Long-read sequencing (LRS), a powerful novel approach, is able to read full-length transcripts and confers a major advantage over the earlier gold standard short-read sequencing in the efficiency of identifying for example polycistronic transcripts and transcript isoforms, including transcript length- and splice variants. In this work, we profile the human cytomegalovirus transcriptome using two third-generation LRS platforms: the Sequel from Pacific BioSciences, and MinION from Oxford Nanopore Technologies. We carried out both cDNA and direct RNA sequencing, and applied the LoRTIA software, developed in our laboratory, for the transcript annotations. This study identified a large number of novel transcript variants, including splice isoforms and transcript start and end site isoforms, as well as putative mRNAs with truncated in-frame ORFs (located within the larger ORFs of the canonical mRNAs), which potentially encode N-terminally truncated polypeptides. Our work also disclosed a highly complex meshwork of transcriptional read-throughs and overlaps.

## INTRODUCTION

Next-generation short-read sequencing (SRS) platforms have revolutionized genomics and transcriptomics sciences, and while they are still invaluable in sequencing studies, the now state-of-the art long-read sequencing methods (LRS) are becoming more popular and represent an even more powerful approach in transcriptome research. Most genes encode multiple transcript isoforms^1^ that are mRNAs or non-coding RNAs (ncRNAs) transcribed from the same locus, but have different transcriptional start sites (TSSs), or transcriptional end sites (TESs), or are the results of alternative splicing^2,3^. The reconstruction of all transcribed isoforms for each gene is challenging with the currently available bioinformatics tools since they have been developed for the analysis of SRS data^4,5^.

Since LRS technologies are able to read full-length RNA molecules, they offer a solution to disclose the full spectrum of complex transcriptomes and offer an insight that is unachievable via SRS methods^6^. LRS platforms are currently commercially available by Pacific Biosciences (PacBio) and Oxford Nanopore Technologies (ONT), which provide read lengths of ~15 kb for PacBio and > 30 kb for ONT that surpass lengths of most transcripts. Both techniques were applied for the investigation of transcriptomic complexity of human cell lines^7^ and various organisms, such as mammals^8^, fish^9^ and plants^10^ and a number of viruses, such as poxviruses^11^, baculoviruses^12^, coronaviruses^13^, circoviruses^14^, adenoviruses^15^; and herpesviruses^16-19^.

Since viral genomes are small and compact, they are ideal subjects for transcriptome analysis with the LRS techniques, as these methods still have a relatively low throughput compared to the SRS techniques^20^. These LRS-based studies repeatedly concluded that transcriptional complexity had previously been underestimated in all of the examined viruses^21^. In addition, ONT is capable of sequencing not only DNA^22^ but also RNA in its native form^23^. Direct RNA sequencing (dRNA-Seq) does not require reverse transcription and PCR amplification therefore, it does not produce spurious transcripts, which are common artifacts of these techniques. While dRNA-Seq has its own limitations^24^, it can be used to validate and to expand cDNA-based LRS studies^16^.

*Human cytomegalovirus* (HCMV, also termed *Human betaherpesvirus 5*) infects 60-90% of the population worldwide^25^ and can cause mononucleosis-like symptoms in adults^26^, and severe life-threatening infections in newborns^27^. It can infect various human cells, including fibroblasts, epithelial cells, endothelial cells, smooth muscle cells, and monocytes^28^. HCMV has a linear double-stranded DNA genome (235 ± 1.9 kbps), which is the largest genome among human herpesviruses^29^. Its E-type genome structure consists of two large domains: the unique long (UL) and the unique short (US), each flanked by terminal (TRL and TRS) and internal (IRL and IRS) inverted repeats^30^. In addition, it encodes four major long non-coding RNAs (lncRNAs) (RNA1.2, RNA2.7, RNA4.9, and RNA5.0)^31^, as well as at least 16 pre-miRNAs and 26 mature miRNAs^32–34^. Although the functions of most genes in infective stages have been identified, many remain uncharacterized^29^. The HCMV genome was shown to express more than 751 translated open reading frames (ORFs)^35,36^, although most of them are very short and located upstream of the canonical ORFs. The compact genome with high gene density has many overlapping transcriptional units, which share common 5’ or 3’ ends, complex splicing patterns, antisense transcription, and transcription of lncRNAs and micro RNAs (miRNAs)^37^. Nested genes are special forms of the 3’-coterminal transcripts, since they have truncated in-frame open reading frames (ORFs), which possess different initiation but have common termination sites^38^. These add even more complexity to the genome regulation and expand coding potential of the virus. Short-read RNA sequencing studies have discovered splice junctions and ncRNAs^39^ and have shown that the most abundant HCMV transcripts are similarly expressed in different cell types^10^.

In our previous work^40^, we used the Pacific Biosciences RSII sequencing platform to investigate the HCMV transcriptome and detected 291 previously undescribed or only partially annotated transcript isoforms, including polycistronic (PC) RNAs and also transcriptional overlaps. However, the RSII method is biased toward cDNA sizes between 1 and 2 kbp, therefore the short and the very long transcripts have not been detected by this analysis. As it was concluded by others^41^, involving other sequencing technologies for the analysis could provide additional insights into the operation of transcriptional machineries of HCMV. Following this concept, in this work, we analyzed the HCMV transcriptome applying a multi-technique approach including ONT MinION and the PacBio Sequel platforms and using both cDNA and native RNA sequencings. Our primary objective was to construct the most comprehensive HCMV transcriptome atlas currently available using data provided by the state-of-the-art LRS methods, and thus to gain a deeper understanding of this important human pathogenic virus.

## RESULTS

### Long-read sequencing of HCMV transcriptome using a multi-technique approach

In this work, we analyzed the HCMV transcripts with ONT MinION technique using cDNA, Cap-selected cDNA, and native RNA libraries and with PacBio Sequel platform using a cDNA library^42^. We also included the data obtained in our previous work using PacBio RSII method^40^. These reads, alongside Sequel and ONT reads, were remapped with minimap2 and reanalyzed with the LoRTIA program developed in our laboratory^43,44^. Figure 1 shows the sequencing platforms, library preparation methods and data analysis steps that were carried out in this work.

**Figure 1.**
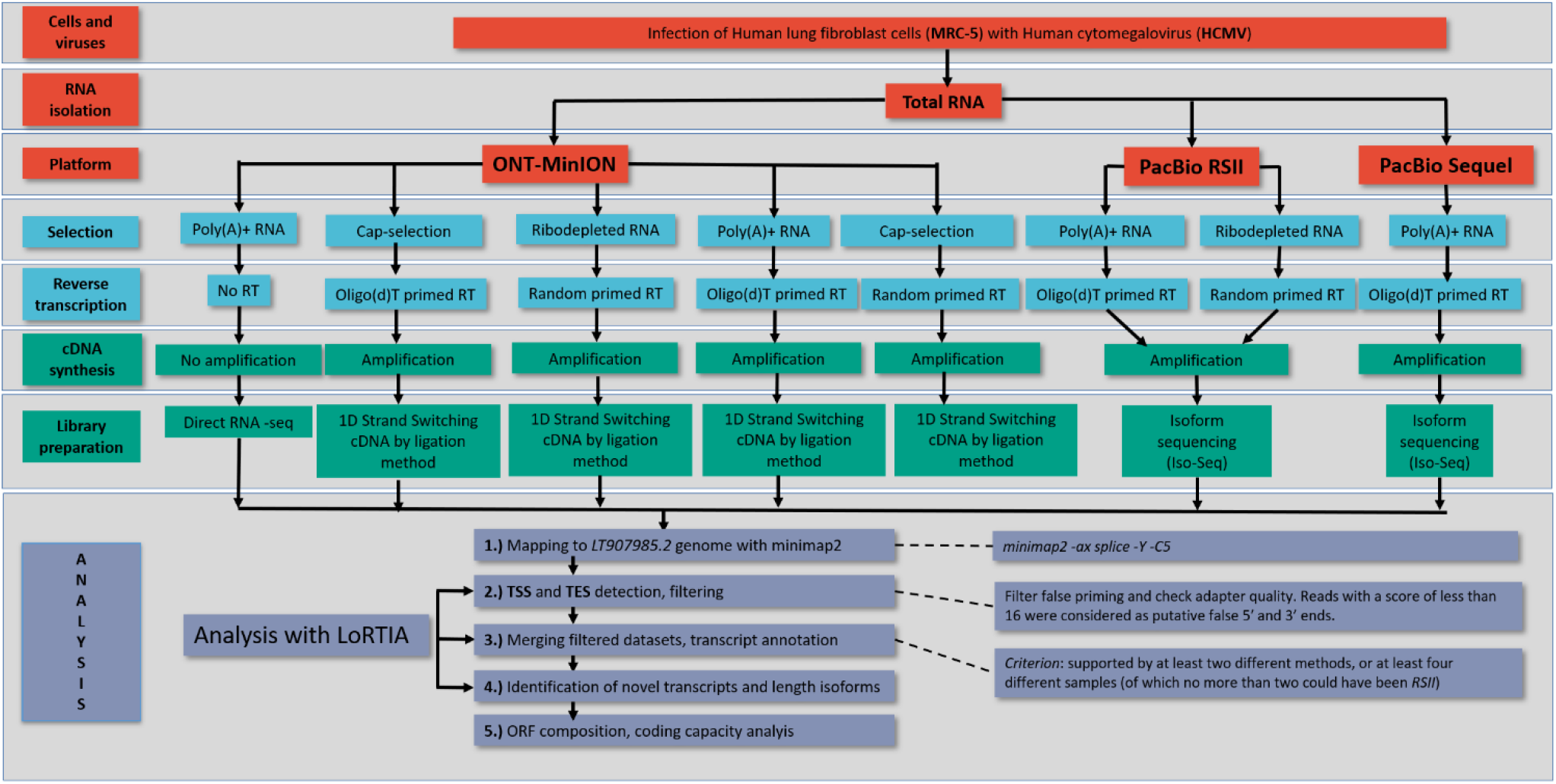
An overview of the utilized sequencing platforms, library preparation and sequencing methods; and bioinformatic analyses of the resulting sequencing reads.

The read statistics of the different sequencing approaches is shown in Table 1, and the read length distributions are illustrated in Figure 2. As the various methods have distinct advantages and limitations, the use of multiplatform transcriptomics approaches have proven to be valuable^43^. For example, dRNA sequencing produces incomplete reads since a 15-30 nt long sequence always lacks from the 5’-termini, and also in many cases poly(A) tails are also missing. Nonetheless, as dRNA-Seq is free of RT- and PCR-biases, it can be used for the validation of introns. However, due to its lower coverage and shorter average read lengths (Table 1 and Figure 1) compared to the cDNA libraries, and its other biases, dRNA-Seq is advised to use in conjunction with other methods. The Cap-selected cDNA library produced the highest throughput, but shorter average read-length due to the applied size-selection (>500 nt). On the other hand, the Sequel library produced the longest reads on average but with a relatively low throughput compared to the ONT cDNA libraries.

**Figure 2.**
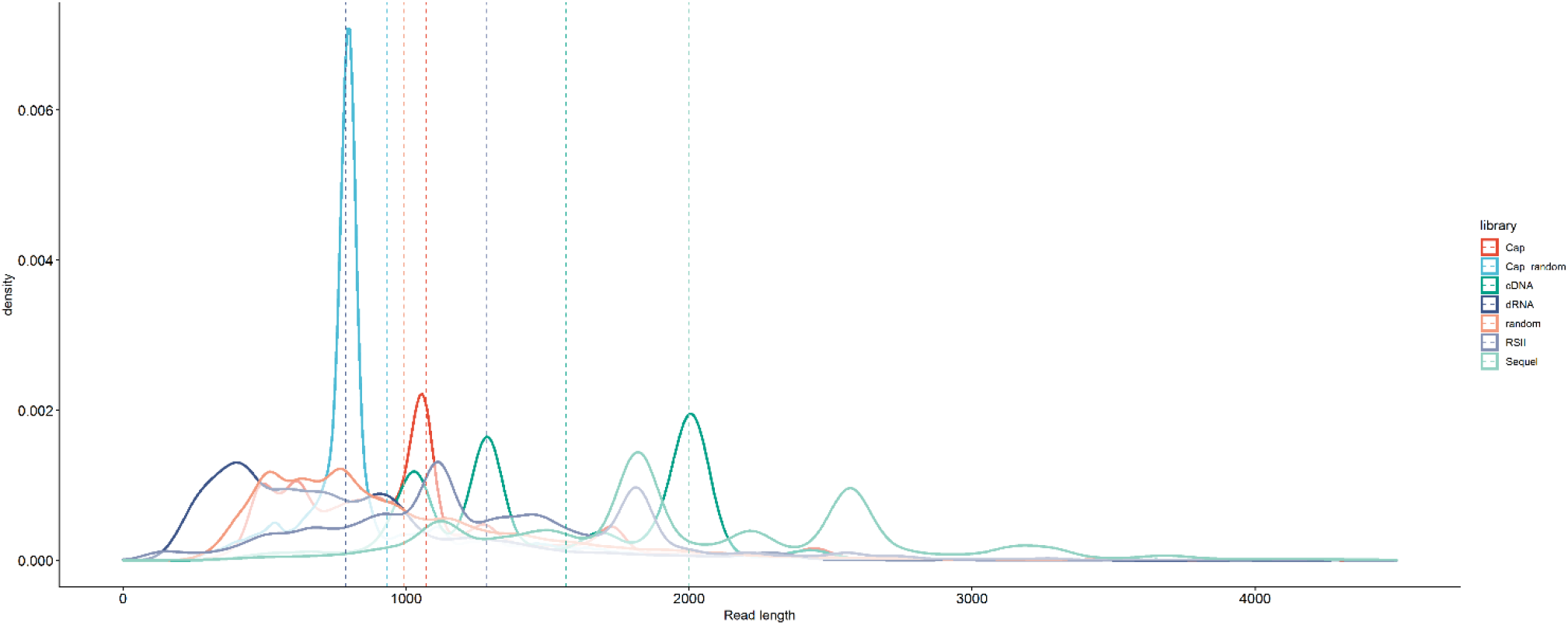
Mapped read densities per sample, with median lengths represented as dotted lines.

**Table 1.** Statistics of the reads, mapped to the Towne varS genome, according to each sample library used.

| *Mapped read statistics* |  |  |  |  |  |  |
| --- | --- | --- | --- | --- | --- | --- |
| *Sample* | *Sample nr.* | *Read count* | *Min. length* | *Mean length* | *Max. length* | *Mean coverage* |
| **RSII** | 8 | 61,375 | 83 | 1,250 | 5,833 | 329 |
| **dRNA** | 1 | 30,366 | 97 | 787 | 8,397 | 101,8 |
| **Sequel** | 1 | 40,233 | 84 | 2,021 | 7,239 | 325,6 |
| **Non-cap oligo(d)T** | 1 | 352,485 | 135 | 1,570 | 10,731 | 2,271.4 |
| **Cap oligo(d)T** | 1 | 576,557 | 118 | 1,073 | 5,137 | 2,245.4 |
| **Cap random** | 1 | 39,776 | 232 | 936 | 8,992 | 159,6 |
| **Non-cap random** | 1 | 151,234 | 190 | 996 | 8,246 | 645,9 |

The LoRTIA software was used to detect the TESs, TSSs and splice sites (hereafter referred to as ‘features’). This program is also able to check the quality of sequencing adapters and poly(A) sequences and to identify and filter out false TESs, TSSs and splice sites generated by RT, PCR and sequencing as a result of false-priming, template switching, RNA degradation^44^, etc. In order to have higher confidence for the validity of the annotated features, stringent filtering criteria were used. The features were only accepted if either one of the two following criteria were met: they were detected by at least two different methods, or they were detected in at least four different samples, of which no more than two could have been RSII samples. For intron annotation, an additional criterion was that every intron had to be supported by at least one read in the native RNA library. In each sample a feature was considered to exist if at least two reads supported it. Subsequently, the LoRTIA software was used to assemble the transcripts based on these features.

The LoRTIA toolkit is able to combine the results of different datasets, therefore this workflow can be regarded as a robust approach for the identification of TESs, TSSs and introns. As a result of the feature detection and subsequent filtering procedure, 93 novel TSSs and 22 TESs were identified. The stringent filtering criteria led to the detection of 103 introns (3 of them are novel), all of them complying with the GT/AG rule. The features that passed the stringent filtering criteria, are termed as ‘validated’ hereafter.

The sequencing libraries were downsampled to the library size of the smallest library (dRNA) in order to be able to compare the efficiency of the different sequencing methods and library preparations in terms of TES, TSS and intron detection. The extent to which the validated features were detected in the downsampled sets show how efficient the respective methods are in terms detecting the TESs, TSSs and introns, regardless of read count and library size. Figure 3 shows the precision and accuracy (recall) of the different libraries in terms of detecting these validated features and the effect of downsampling. In terms of intron detection, the highest recall was achieved in the Sequel sequencing library (71%), which was comparable to that of the dRNA library. RSII showed a somewhat lower recall, but its precision was somewhat higher. With the exception of the Cap-selected cDNA sample, which was similar to the PacBio samples, the precision was generally higher in the ONT samples (less false positives), while the recall was lower (less true positives). In the case of full libraries, the poly(A)-selected Cap sample showed a very high recall (95%), but many false positives as well, as this library was the largest; while the random Cap sample a high precision, but there were many valid features that it could not detect (recall=22%), again because of the library size, which in this case was small, comparable to that of the size of the dRNA sample. TES detection was more efficient in the PacBio samples, as both the precision and recall were higher in the downsampled libraries, (except for the dRNA and the poly(A)-selected Cap samples whose precision was similar, but not their recall). Moreover, the full PacBio libraries showed comparable values to the full ONT libraries, despite their much lower coverage. The Sequel sample performed the best (94% recall and 81% precision), even though its read count was an order of magnitude lower than that of the Cap and non-Cap cDNA samples (Table 1). In the case of TSSs, the PacBio samples showed a better performance as well than the Cap and non-Cap cDNA samples, both in the downsampled and full libraries. Thus, generally, the PacBio samples showed better efficiency in terms of TES and TSS detection than the ONT libraries, and similar performance in intron detection to the dRNA library; as after downsampling, both their precision and recall values were higher.

**Figure 3.**
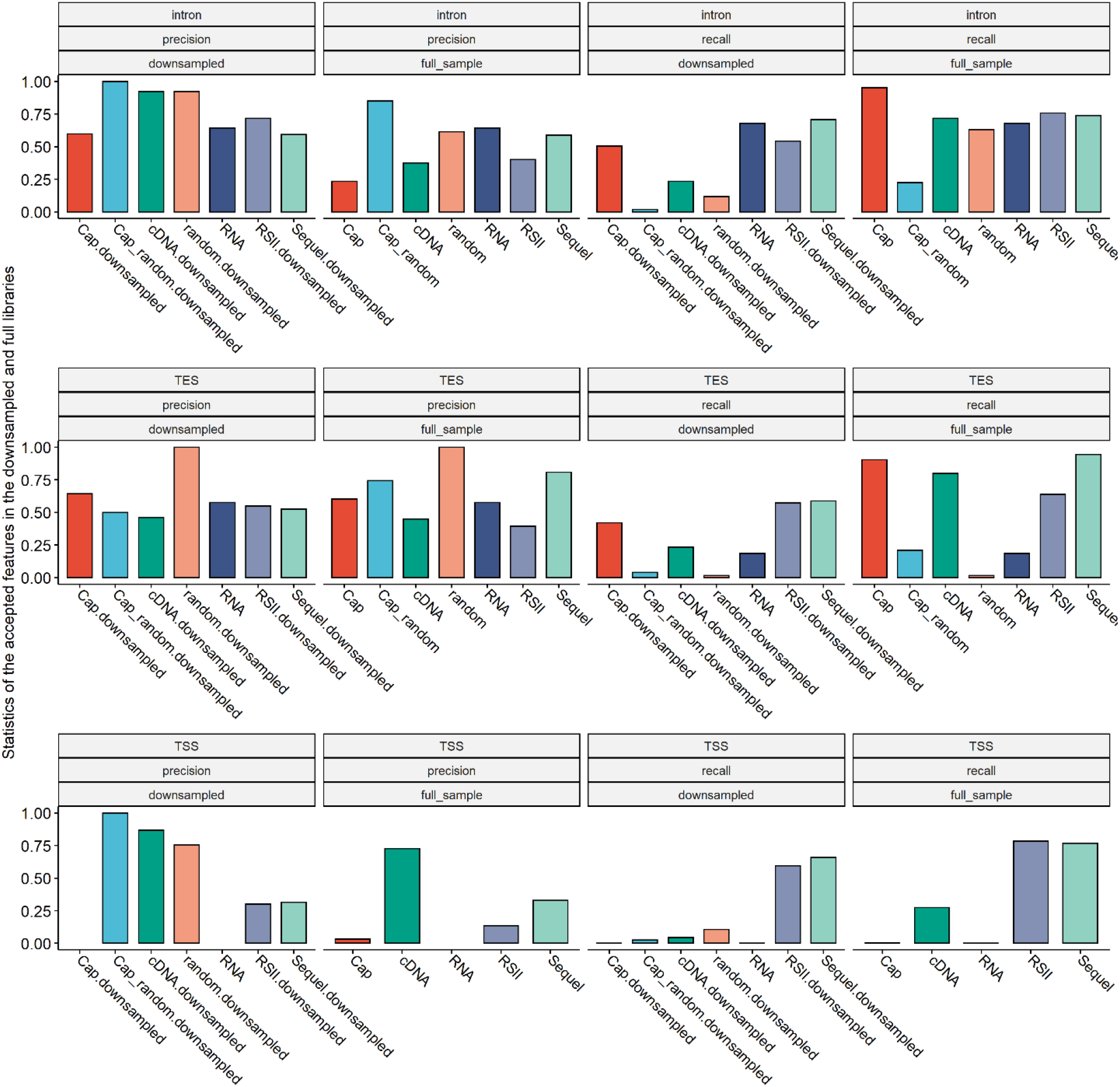
Effect of downsampling on the recall and precision of the detection of the validated features (introns, TESs and TSSs) in the different sequencing methods and library preparations. The features were termed validated if they passed the stringent filtering criteria. Recall was calculated as the ratio of validated features that was found in the respective samples, while precision was calculated as the ratio of true positives in all the hits of respective samples. The downsampling of reads were carried out to match the sizes of the libraries to the size of the smallest library (dRNA).

Next, we used the *transcript_annotator* function of LoRTIA with the filtered feature sets (TESs, TSSs and introns) for the annotation of transcripts. Then, we compared the identified transcripts with the previous dataset^40^ and with other literature sources (Supplementary file S2). We termed a transcript identical to another if their termini were within a 10-nt window, and if their intron composition matched. If the difference was larger than 10 nt or the intron composition differed, then the transcript was termed either a length isoform or a splice-variant, respectively. Categorizing and naming of transcripts were carried out in-line with our previous convention^45^.

After the stringent filtering procedure, a total of 437 transcripts were annotated (Figure 4). This is a significant increase compared to the previously described 291 transcripts using RSII sequencing, highlighting the advantages of using various sequencing methods. Although, 242 transcripts have already been described: 183 in our previous dataset^40^ and 59 in other sources, the remaining 195 transcripts are novel. Note that the lack of confirmation of certain earlier described TSSs, TESs, introns and transcripts due to using more stringent criteria for the annotation in this study, does not necessarily mean that they do not exist.

**Figure 4.**
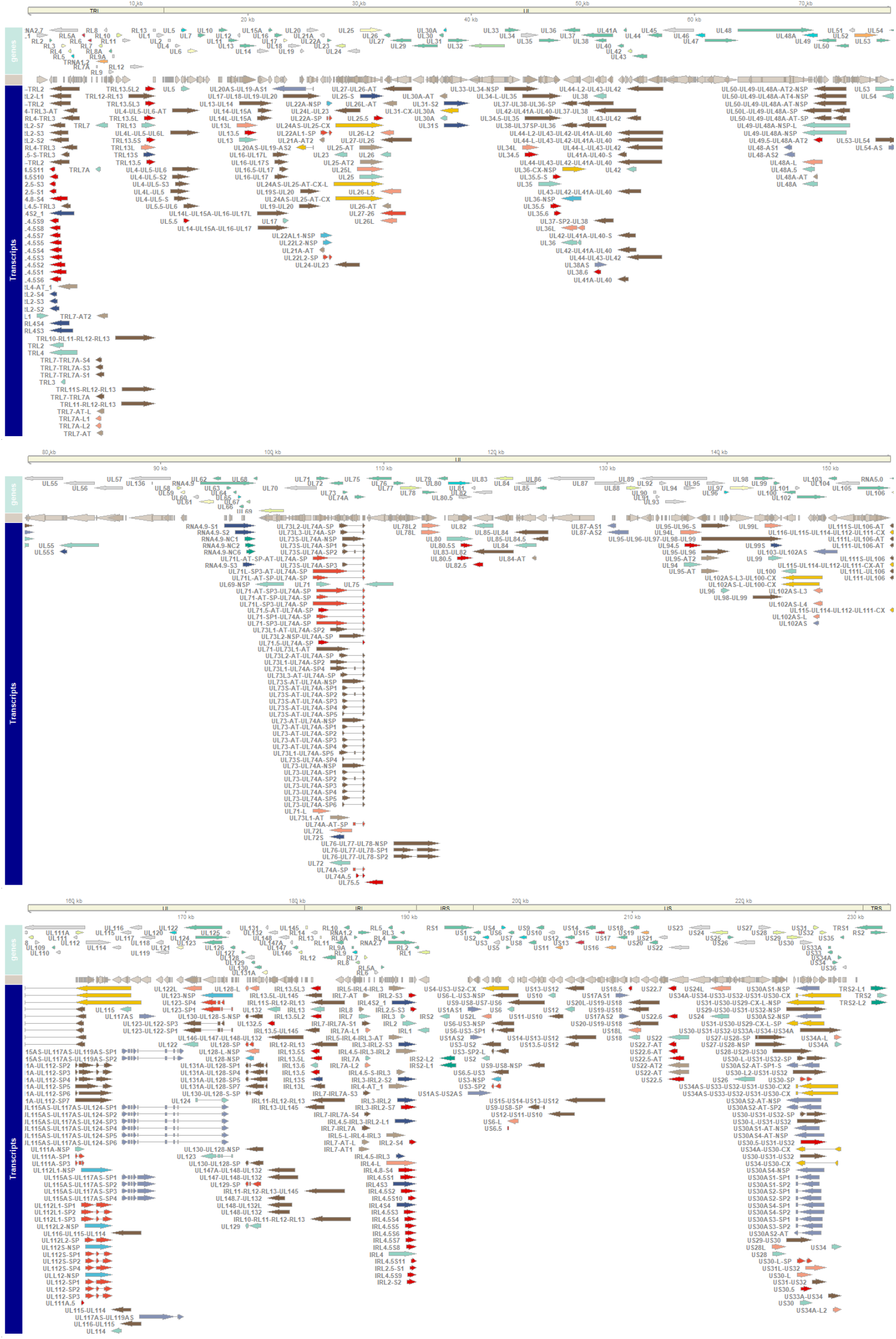
Genes (upper panel), ORFs (middle panel) and the annotated transcripts (bottom panel) of HCVM LT907985.2. Genes are colored according to their functional classes, as described in^29^, while transcripts are colored according to their category.

### Transcript isoforms

The alternative use of transcription start sites (TSS isoforms) results in a shorter (labeled here: ‘S’) or longer (‘L’) 5’-untranslated regions (5’-UTRs) compared to the canonical transcript, but containing the same main ORFs, although short upstream ORF (uORF) compositions may differ. In this dataset, we detected 10 novel short TSS isoforms, which include short isoforms of the RL4, RL2 and RL3 transcripts (3, 3 and 2 isoforms, respectively), as well as one isoform of UL31 and UL25 transcripts (Figure 4). We used the reads obtained by dRNA sequencing to validate the TSSs of the short length isoforms. Since dRNA sequencing reads lack 10-25 nucleotides from their 5’-termini (Supplementary Figure S1), we accepted those transcripts, where at least one (LoRTIA filtered) dRNA read mapped (in the correct orientation) near their 5’-ends, and the read was shorter than the transcript but with no more than 25 nucleotides. We also detected 16 novel long TSS isoforms: three from the *rl7A* gene, and two long TSS isoforms *ul102, ul128, ul26,* and *us34* genes, and one from the *rl4, rl7, ul71, ul72* and *us6* genes as well. At the *rl4-rl8* genomic region transcripts use both alternative TESs and TSSs (Figure 4).

In this study, 16 novel alternative TES variants (labelled with–AT in their names) were also identified. More than half of them were produced by the *rl4, rl5* and *rl7* genes (2, 3 and 5 isoforms, respectively). Two TES isoforms of the UL25 and two of the UL95 mRNA were identified as well, and one from both *ul21A* and *us2* genes.

In addition, we detected 8 novel splice isoforms of which one is an intron-retention variant (NSP, non-splice variant) of *ul123.* Three SP isoforms were found from the ul71 gene, while the rest were expressed from the *ul106-129* region: *ul112, ul123* and also and *ul129* genes were found to express novel splice variants (2, 2 and 1, respectively).

### Non-coding transcripts

In this work, 96 RNA molecules were identified that did not fully encompass any of the 385 long ORFs (>10 AA), described via ribosome-profiling by Stern-Ginossar et al^35^. However, many of these transcripts did contain experimentally validated short ORFs from the same source and *in silico* predicted in-frame co-terminal ORFs (some transcripts of rl2-rl13, ul73, us22, ul111A, ul123, ul128, ul129 and ul132), or only predicted ORFs (some transcripts of genes ul82 and us33). UL54.5 is an exception, as the *in silico* predicted ORF that it carries is not co-terminal with the canonical UL54 ORF, but it is in-frame with it. These transcripts are considered *embedded* transcripts rather than non-coding and will be discussed in the following section. The rest of the non-coding transcripts include 3 novel lncRNAs found to be expressed from the non-coding gene *rna4.9,* and 4 from the *rs2* gene.

Another important types of lncRNAs are the antisense RNAs (asRNAs) molecules that are controlled by their own promoters^20,46^. We identified 9 such asRNAs from the *us30* gene region. The only gene in the vicinity that is in the same orientation as these transcripts is the *us33*, which starts 764 nts downstream to the start of these transcripts. It is unlikely however, that its promoter is involved in the transcription of these asRNAs, as no detected transcripts initiated from that gene. In addition, many transcripts contain antisense segments, but they cannot be considered as true AS transcripts, rather they are the products of either transcriptional read-through between convergent genes (as is the case in TES isoforms encoded by *us1, ul48, ul53, ul54, ul8, ul89, ul103, ul115* and *us33*), or transcriptional overlaps between divergently oriented genes (*ul20, ul123* and *us34*).

### Transcriptional overlaps

We carried out a genome-wide analysis of transcriptional overlaps: the number of overlapping transcripts were calculated using a 10-nt sliding window. The transcriptional orientation of genes relative to the adjacent genes can be either convergence (→←), divergence (← →), or co-orientation (→→)^20^. Parallel overlaps between co-oriented genes are common throughout the entire viral genome^38^, whereas divergent and convergent overlaps are restricted into distinct genomic locations (Supplementary File 1. - Supplementary Figure S2). We found two novel convergent transcriptional overlaps, one between transcripts encoded by *us32* and *ul33* genes and another encoded by *ul112* and *ul114* gene pairs. We also detected a novel divergent transcriptional overlap in the *ul100-102* region.

### Multigenic transcripts

Long read sequencing techniques are especially suitable for distinguishing the polycistronic mRNAs from monocistronic transcripts^18^. We detected 90 novel polycistronic (PC) and 15 complex transcripts (CX) (Supplementary Figure S3). CX transcripts are multigenic RNA molecules that contain at least one oppositely oriented gene^49^. 13 PC transcripts contain either additional introns or genes which were earlier considered to produce exclusively monocistronic transcripts or contain novel introns. Novel polycistronism was found in the following genes: *us7, us3, us28, us19, us12, ul71, ul50, ul38, ul34, ul27, ul17 and ul116.* We also identified 12 novel CX transcripts which were generated by transcriptional readthroughs or the use of long 5’-UTRs. The rest of the multigenic transcripts are TSS or TES variants of already described PC or CX RNA molecules. The identified PC transcripts include 48 bicistronic, 22 tricistronic, 12 tetracistronic, 5 pentacistronic, 2 hexacistronic and 1 septacistronic molecules (Supplementary Figure S3). Polycistronic transcripts of nested genes were also found in *ul34* (1) and *ul148* (1).

In the *ul89-99* region, until now the polycistronic transcription units were shown to produce two families of nested 3’-coterminal transcripts, encompassing ORFs UL92-UL94 and UL93-UL99, respectively that are differentially regulated^48^. Here we identified two novel TESs that produce seven novel TES transcripts, two in UL95 and three in UL96, respectively (Figure 4, Supplementary File 2). The transcripts associated with the former TES are partially antisense to UL89 and carry three distinct in-frame ORFs (two short and one long), whereas transcripts associated with the latter TES are either monocistronic transcripts of UL96 or are polycistronic UL95-UL96. The bicistronic UL95-UL96 transcripts carry the same ORFs. Until now, transcripts produced from this region always described to end in UL99 and no data was found on the monocistronic, or bicistronic transcripts. UL95 protein is required for late viral gene expression and, consequently, for viral growth^46^ and is associated with latency^47^, thus their monocistronic variant may be important in regulating late gene expression as well.

Polycistronic transcripts have already been described in the *us6-us11* region^51^, but the US9-US6 transcript is novel. We also discovered a novel polycistronic RNA molecule (US12-US10) in the *us12-17* genomic region^48^.

In addition, six PC transcripts were expressed from the *ul42-44* region, and two from the *ul130-ul132* region; and we also identified several polycistronic transcripts in the RL7-RL3 and in the RL11-RL13 region. A bicistronic IRS2-S2-RL1 transcript, was also identified, which carries the IRL1 ORF and shares its TSS with the IRS2 mRNA. The highest number of novel PC transcripts (17) were found in the *ul71-ul73* region; however, these are length variants of already described PC transcripts.

We detected two complex transcripts of UL101 which are antisense to UL101 and partially antisense to UL102 but sense to UL100. These are probably the products of the downstream convergent gene *ul103.* Antisense transcripts were described already from this gene^53^, however none of them contains the ORFL234C ORF (132 AA), which is located in the intron of the two spliced mRNAs encoded by this gene^53^.

We identified nine PC and two CX transcripts at the *ul112-116* genomic region (Figure 4). The *ul112-113* region encodes four phosphoprotein isoforms, two of them are essential for viral DNA replication^54^, whereas the *ul115-116* region encodes envelope glycoproteins^55^. The complex transcripts in this region are the results of the transcriptional readthrough of the tandem *ul115* and *ul116* genes across the full length of the oppositely oriented *ul112* gene.

### Nested genes

The use of alternative TSSs may result in the expression of 5’-truncated transcripts-that lack canonical ATG start codons. Herein, the canonical ORF contains one or more 5’-truncated in-frame 3’ co-terminal ORF(s), which if functional can be considered as nested protein-coding genes^51^. The embedded ORFs have been analyzed using ribosome-profiling by Stern-Ginossar and coworkers^35^ (referred to as ‘validated’ hereafter). We predicted ORFs on the viral genome (the resulting putative ORFs are termed ‘predicted’ hereafter) and found that 15 truncated transcripts carry novel ORFs. If translated, the truncated mRNAs lead to an N-terminally truncated version of the protein encoded by the canonical ORF. Overall, 63 such transcripts were identified in this sequencing dataset of which 40 were described previously and 23 is novel (each of these were validated by dRNA sequencing). Truncated transcripts for novel putative nested genes were found within the following canonical genes: *rl2, rl3, rl13, ul111A* and *ul48A*. Additional novel truncated transcripts were found in genes, where such transcripts have already been described in genes *ul34, ul148, ul94* and *rl4*.

We carried out a promoter analysis of the transcripts: the presence and sequence composition of CAT, GC and TATA-boxes and their distances from the TSS of the transcripts were examined. In addition, the Kozak consensus sequences of the transcripts were analyzed as well. We then compared these features of the 5’ truncated RNAs (with truncated ORFs) to the transcripts of their host genes, those that carry canonical ORFs. Differences in the sequence compositions of their promoter elements were detected (Figure 5), however the number of transcripts where CAT-boxes were found was only 3 (both among the truncated and the canonical transcripts). GC-boxes and TATA-boxes were found in more cases (Figure 5) and an apparent difference was seen between the two transcripts groups, indicating that the transcriptional regulation of the embedded genes differ from their hosts. TSSs showed clear differences as well: the bases upstream of the C/G start site contain more A-s in the truncated transcripts (Supplementary Figure S4 A). The Kozak sequence composition showed a modification mainly in the important −3 Kozak site from the consensus G/A to T/G, which weakens the translation initiation signal somewhat, however this did not cause an overall significant decrease of the mean Kozak sequence score in these transcripts (Supplementary Figures S4 B and C). The differences altogether suggest that besides coding for different protein products, the embedded genes are differentially regulated both on the translation and on the transcription level, compared to their canonical counterparts.

**Figure 5.**
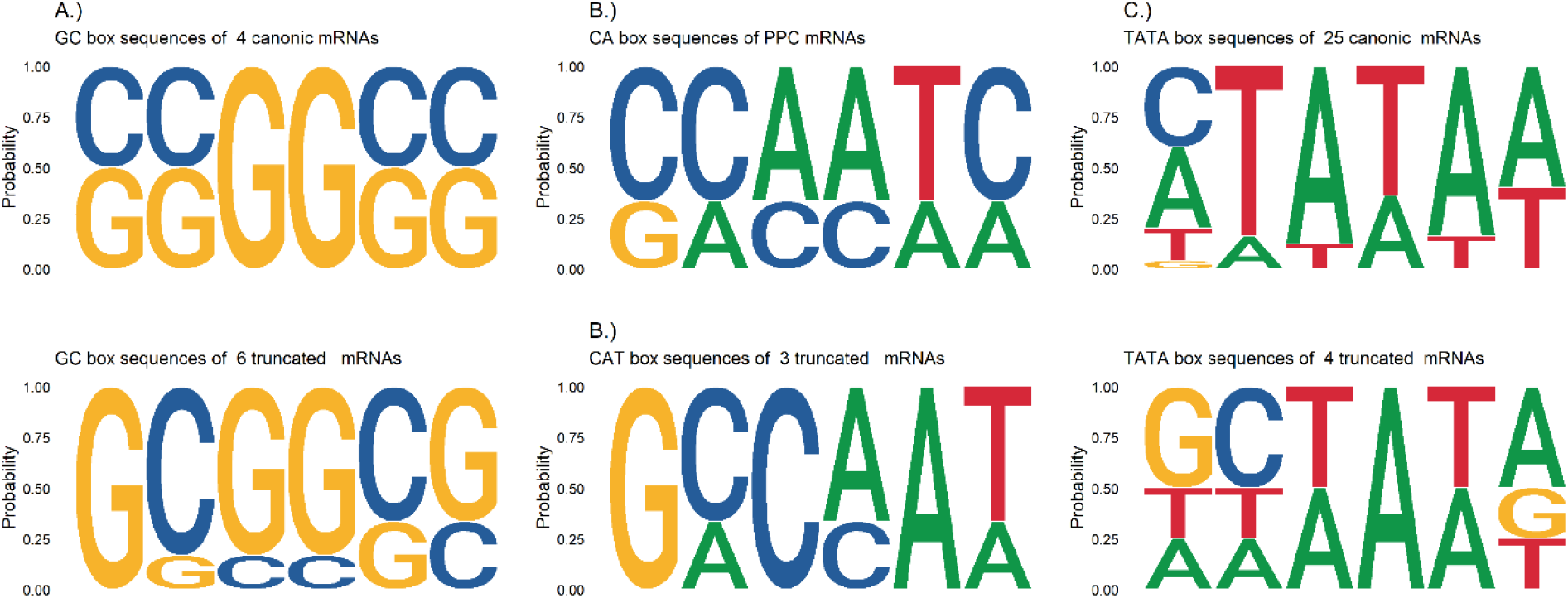
Weblogo of promoter elements: GC-boxes A.), CAT-boxes B.) and TATA-boxes C.) of truncated and canonical transcripts. Only those host genes were selected for the comparison that contained embedded genes. The number of each transcript that was found to contain promoter elements are shown above the respective weblogos.

### uORFs

Upstream ORFs are a class of cis-regulatory elements within the 5’ UTR of the respective mRNAs, and represent an alternative mode of translational regulation^57^. The uORFs are short (<30 codons) and may initiate at near-cognate start codons; they generally repress the translation of the downstream main ORF, however they can stimulate translation of the downstream coding region as well^58^. The uORFs composition in transcript variants (e.g. in short or long TSS variants) encoded by the same gene modulates their translational regulation^59^. In order to assess the coding capacity of the HCMV transcriptome, we transferred the experimentally validated ORF list (both short and long) published by Stern-Ginossar and collegues^35^ to the Towne varS genome (from the Merlin genome) using BLAST. We included those ORFs from the *in silico* predicted list that were co-terminal and in-frame with the canonical ORFs (these are some of the embedded genes of the truncated transcripts) to the resulting ORF list. Subsequently, we mapped them to the detected transcripts and compared these ORF compositions to what was previously described^40^. Figure 6 upper panel shows the distribution of these ORFs in the transcripts and which dataset that are derived from. The analysis revealed 149 novel combinations of the validated ORFs (considering both short and long) carried by the transcripts. This is a ~38% increase compared to the previously described 247, which greatly expands our earlier estimate of the coding capacity of the HCMV genome.

**Figure 6.**
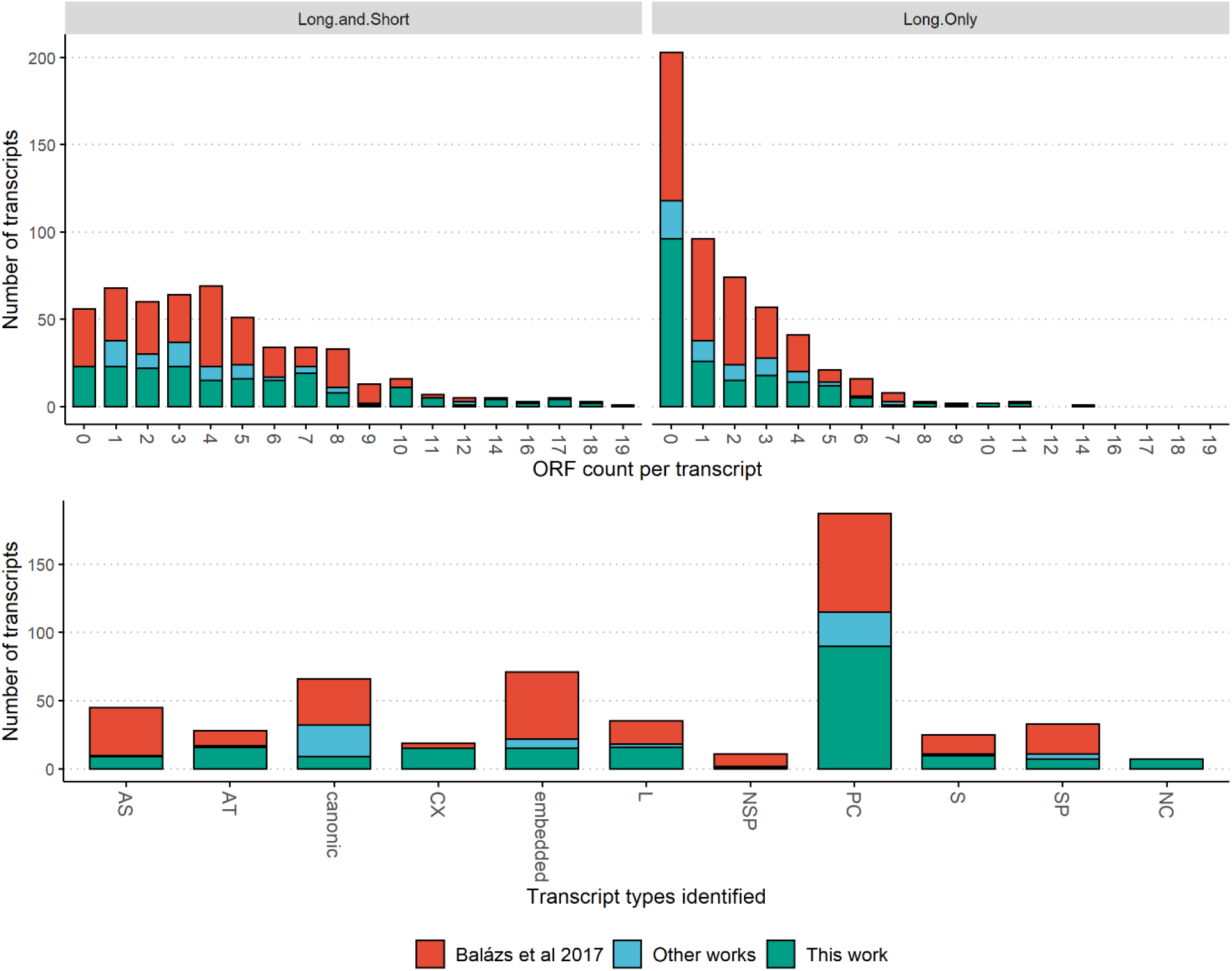
Upper panels: The frequency of transcripts, according to the number of ORFs they encompass (Long, or Long+Short, according to^35^). Lower panel: number of transcripts identified according to transcript categories. Transcripts described in this work are colored with blue, those that we described previously^40^ are colored red, and those that we detected but were described in other works previously are colored green.

## DISCUSSION

In this work, we applied two LRS platforms (ONT and PacBio) with various library preparations and sequencing methods: native RNA and cDNA with and without Cap-selection and using oligo(d)T, or random primers to re-annotate the HCMV transcriptome. We used minimap2 for mapping and the LoRTIA software, with a stringent filtering procedure, to identify transcripts, which lead to the identification of 26 novel TSSs, 34 novel TESs and 25 novel introns. Comparison of the downsampled libraries showed differences between the library preparation protocols and sequencing methods: generally, the PacBio samples showed better performance than the cDNA ONT libraries in terms of TES and TSS detection, and similar performance to that of the dRNA library with respect of intron detection.

We then used the identified features to assemble HCMV transcripts, which enabled the confirmation of 242 previously described, and the identification of 195 novel transcripts. The novel transcripts include 7 lncRNAs, 15 complex, 9 antisense, 23 putative protein coding (truncated), 90 polycistronic, 7 splice variant, 1 intron-retention variant, 16 5’-long 10 5’-short variants and 16 alternatively terminated transcripts. The novel multigenic transcripts include 65 PC transcripts that are expressed from genes where polycistronism has not been described before or contain novel introns.

Many transcript isoforms were produced from *rl2-rl7* genes. A previous report detected a high transcriptional activity from the RL genes, mainly *rl4*^39^. The *ul71-ul73* region expressed many length variants of previously described PC transcripts.

In the *ul89-99* genomic region, we identified novel bicistronic and monocistronic transcripts that are likely be differentially regulated as is the case with their the previously identified variants^48^. Although UL95p is required for viral growth, probably these variants were downregulated in previous studies to such an extent that they have gone undetected. In the *us30-34* region two PC, seven CX, and 14 AS transcripts were detected.

We also identified two complex transcripts of UL102, which are antisense to UL101 and partially antisense to UL102 but sense to UL100. Previous studies confirmed antisense transcripts from this gene^53^, however none of them contains the ORFL234C. We also identified two PC and three CX transcripts from the UL115-UL116 region. At this point we can only speculate about the function of the complex transcripts. It is possible that the overlap causes transcriptional interference, as proposed earlier^18,60^.

Alternative TSS-usage can cause the 5’-truncation of the ORFs, which may result in the formation nested genes, wherein one or more ORFs that are in-frame and co-terminal with the canonical ORF. These putative embedded genes might encode N-terminally truncated polypeptides. The identification of novel TSSs enabled us to confirm the existence of many such embedded genes. To gain high confidence in the existence of their expressed truncated mRNAs, an additional validation step (using dRNA reads) was employed to the already stringent transcript filtering criteria. These truncated mRNAs represent potentially novel functionalities of the viral protein repertoire, if translated. The promoters of nested genes and their sequence composition around the TSSs differ substantially, compared to the host genes, which may refer differences in the transcriptional regulation. Their Kozak sequences also show dissimilarities, thus they are presumably regulated differentially on a translational level as well.

We found that the detected nested genes and their truncated transcripts, along with multigenic transcripts and transcript isoforms carrying different ORF compositions (many times attributed to different uORF compositions) significantly expand the coding capacity of HCMV. The previously estimated number of unique ORF composition carried by the transcripts was increased with 38%.

Human Cytomegalovirus is a highly prevalent infectious agent, which partly due to its complex genome and transcriptional architecture causes a lifelong infection. By using novel RNA sequencing methods, a deeper insight into its intricate transcription was achieved in this study. The herein reported results complemented with the previous ones represent the most detailed transcriptome of this important virus.

## MATERIALS AND METHODS

In this article we used the sequencing datasets described in^52^ for Biosample ERS1870077 and ERS2312967 and^42^ for ERS2312967. The materials and methods used to generate those data are described here as well.

### Cells and viruses

Human lung fibroblast cells [MRC-5; American Type Culture Collection (ATCC)] were cultured at 37°C and 5% CO2-concentration in DMEM supplemented with 10% fetal bovine serum (Gibco Invitrogen), 100 units of potassium penicillin and 100 μg of streptomycin sulfate per 1 ml (Lonza). Four and eight T75 cell culture flasks were used for the ERS2312967 and ERS1870077 samples, respectively. Semi-confluent cells were infected with the Towne VarS (ATCC) strain of HCMV (a multiplicity of infection of 0.5 plaque-forming units (pfu) per cell). The viral-infected cells were then incubated for 1 h, which was followed by the removal of the culture medium and washing of the cells with phosphate-buffered saline. Subsequently, fresh culture medium was added, and cells were kept in the CO2 incubator for 24, 72, or 120 h, in case of ERS2312967, and for 1, 3, 6, 12, 24, 72, 96 or 120 h in case of ERS1870077.

### RNA purification

Total RNA was extracted from each time point samples using the NucleoSpin RNA kit (Macherey-Nagel) as described in our earlier publications (Oláh et al, 2015 BMC Microbiology, Tombácz et al 2018 Sci Data PRV).

### PolyA(+) RNA purification and ribosomal RNA removal

20 μl of the isolated RNA sample from each time point were pooled. The Oligotex mRNA Mini Kit (Qiagen) was used to select polyadenylated RNAs from both samples. Two different, poly(A)-selected libraries were prepared. For the analysis of the non-polyadenylated RNA fraction of the viral transcriptome, the ribosomal RNAs were removed using the RiboMinus™ Eukaryote System v2 (Thermo Fisher Scientific) according to the kit’s instructions.

### Library preparation and sequencing

#### Biosample ERS2312967

##### ONT MinION sequencing – direct RNA

500 ng polyA-selected RNA was used for direct RNA sequencing. First-strand cDNA was generated using SuperScript IV (Thermo Fischer Scientific) and the adapter primers (supplied by the ONT’s Direct RNA Sequencing kit; SQK-RNA001). The library preparation was carried out with the ONT Ligation Sequencing 1D kit (SQK-LSK108) following the recommendations of the manufacturer.

##### ONT MinION sequencing — oligo(dT)-primed, Cap-selected cDNA

Two micrograms from the total RNA sample was used to generate first strand cDNAs using the Lexogen TeloPrime Full-Length cDNA Amplification Kit. Oligo(dT) or random primers were used for the reverse transcription (RT). The ligation of the 5’ adapter to the samples was carried out overnight at 25°C. The samples were amplified with PCR (30 cycles) using the reagents supplied by the TeloPrime kit. The libraries for nanopore sequencing were generated using the Ligation Sequencing 1D kit (SQK-LSK108, ONT) and the NEBNext End repair / dA-tailing Module NEB Blunt/TA Ligase Master Mix (New England Biolabs) according to the manufacturers’ instructions.

##### ONT MinION sequencing - random-primed, non-Cap-selected cDNA

RNA mixture from the rRNA-depleted sample was used to produce cDNA library for MinION sequencing. The RT reaction was carried out according to the ONT’s Ligation Sequencing 1D kit (SQK-LSK108) using random primers instead of oligo(dT) primers and SuperScript IV (Thermo Fischer Scientific). The second-cDNA strand was primed with the strand-switching (5’) adapter. The amplification of the samples was carried out with KapaHiFi DNA polymerase (Kapa Biosystems) enzyme applying 16 PCR cycles. For the ligation reaction, the NEBNext End repair / dA-tailing Module NEB Blunt/TA Ligase Master Mix (New England Biolabs) was used.

##### ONT MinION sequencing - random-primed, Cap-selected cDNA

The Lexogen TeloPrime Kit was used to generate libraries from 5’-Capped RNAs with random primers to enrich the non-poly(A)-tailed RNAs or to capture the 5’-end of rare, very-long complex transcripts. The random-primed RT was followed by the ligation of the 5’ adapter from the TeloPrime Kit at 25°C, overnight. The sample was amplified through 30 PCR cycles with the KapaHiFi DNA polymerase enzyme (Kapa Biosystems). The library for nanopore sequencing was generated using the ONT’s Ligation Sequencing 1D kit (SQK-LSK108) according to the manufacturer’s recommendations.

All the ONT libraries were run on R9.4 SpotON Flow Cells with a MinION sequencing device.

#### Biosample ERS1870077

##### PacBio RSII sequencing

A total RNA mixture containing samples in equal volume from all post infection (p.i.) time points was used for library preparation. For the enrichment of polyA-tailed RNAs, the Oligotex mRNA Mini Kit (Qiagen) was used. The Eukaryote System v2 (Ambion) kit was utilized to produce ribosomal RNA-free samples for random primer-based sequencing. Adapter-linked anchored oligo(dT) primers or random primers were used for the RT. The cDNA samples were prepared using the Clontech SMARTer PCR cDNA Synthesis Kit. The reactions were carried out according to the PacBio Isoform Sequencing (Iso-Seq) protocol. PCR reaction was performed on 16 cycles and 500 ng from the amplified samples were used to prepare the PacBio SMRTbell libraries with the PacBio DNA Template Prep Kit 2.0 and they were bound to MagBeads (MagBead Kit v2). The P6-C4 chemistry was used for sequencing. RSII SMRT Cells (v3) were applied with the RSII platform. Seven SMRT cells were utilized for sequencing the poly(A)+ SMRTbell templates and one for the random-primed library.

##### PacBio Sequel sequencing

RNA mixture from the polyA(+) samples was used to generate cDNA library for single-molecular real-time sequencing on PacBio’s Sequel platform. The Clontech SMARTer PCR kit and the PacBio Iso-Seq protocol were used. The SMRTbell DNA Template Prep Kit 2.0 was used for the generation of libraries, which then were bound to Sequel DNA Polymerase 2.0. The PacBio’s MagBead-loading protocol was used with MagBead Kit 2.0, and the sequencing was carried out on the Sequel instrument using a single Sequel SMRT Cell (v2) 1M with Sequel Sequencing chemistry 2.1. The movie length was 10 h. Consensus sequences were generated with the SMRT-Link v5.0.1 software (Potter, 2016).

The optimal conditions for primer annealing and polymerase binding were determined with the PacBio’s Binding Calculator in RS Remote.

##### ONT MinION sequencing – oligo(dT)-primed, non-Cap-selected

PolyA(+) RNA fraction was used to generate cDNAs using SuperScript IV (Thermo Fischer Scientific) and adapter-linked oligod(T) primers. The 5’ adapter primers were ligated to allow for second-strand synthesis. The sample was amplified through 16 cycles using KapaHiFi enzyme. The PCR products were run on an UltraPure Agarose (Thermo Fischer Scientific) gel and cDNA fragments larger than 500 nt were purified using the Zymoclean Large Fragment DNA Recovery Kit (Zymo Research). The library was prepared using the ONT 1D kit (SQK-LSK108) and the NEBNext End repair / dA-tailing Module NEB Blunt/TA Ligase Master Mix (New England Biolabs) according to the kit’s manual. The library was sequences on a R9.4 SpotON Flow Cells using a MinION device.

### Data Validation

The measurement of the samples was carried out with Qubit 2.0 fluorometer (Life Technologies) while their quality was checked by Agilent 2100 Bioanalyzer (Agilent High Sensitivity DNA Kit). Samples with RNA Integrity Numbers higher than 9.5 were used for this study.

### Read Processing

All sequencing reads were aligned to the HCMV strain Towne VarS genome (LT907985.2) using minimap2^62^, using options *-ax splice -Y -C5*. The mapped reads were not trimmed and may therefore contain terminal poly(A) sequences, 3’ adapters or 5’ adapter sequences (AGAGTACATGGG in case of the Sequel, TGGATTGATATGTAATACGACTCACTATAG in the case of the CapSeq and TGCCATTACGGCCGGG in case of the not cap-selected cDNA sequencing). These sequences are soft clipped and can be used to determine read strandedness. Direct RNA sequencing reads do not contain 5’ adapters; read directions are determined by the sequencer as RNA molecules enter the nanopores with the poly(A)-tail first.

Read statistics were calculated using custom R scripts (available upon request).

### Feature detection and transcript annotations

The LoRTIA toolkit (https://github.com/zsolt-balazs/LoRTIA), was used with default parameters for the both the PacBio (*LoRTIA –5 AGAGTACATGGG --five_score 16 --check_in_soft 15 –3 AAAAAAAAAAAAAAA --three_score 18 -s poisson -f True*) and ONT (*LoRTIA –5 TGCCATTAGGCCGGG --five_score 16 --check_in_soft 15 –3 AAAAAAAAAAAAAAA --three_score 16 –s poisson -f True*) mapping outputs to annotate introns, TSSs, and TESs and the results were combined with the *sum_gffs.py* command. In order to make sure that these features are consequently valid, they were only accepted if either one of the two following criteria were met: a given feature was detected by at least two different methods, or it was detected in at least four different samples, of which no more than two could have been RSII samples. For intron annotation, we set an additional criterion: one of the samples that supported it had to be the native RNA library.

Transcript naming scheme: we used the same transcript naming scheme as in our previous studies^45^. Genome annotations (including ORFs, according to Stern-Ginossar et al., 2012^35^) were transferred to the Towne var-S genome with metablastr^63^ and Liftoff ^54^.

Data evaluation, comparison of the transcripts, assessing their coding potentials, calculating their Kozak sequence score, carrying out ORF predictions and BLAST comparisons and generating visualizations were carried out with the following R packages: ORFik^65^, Gviz^55^ and tidygenomics^58^ using custom R scripts. Promoter elements and PASs were searched with https://github.com/moldovannorbert/seqtools.

## Data Availability Statement

Raw and mapped data files have been uploaded to the European Nucleotide Archive under the accession number PRJEB25680 (https://www.ebi.ac.uk/ena/data/view/PRJEB25680) and PRJEB22072 (https://www.ebi.ac.uk/ena/data/view/PRJEB22072). All data can be used without restrictions.

## AUTHOR CONTRIBUTIONS

DT and ZC performed the sequencing work and the library preparations. KM cultured the cells and carried out the infections. BK wrote the scripts and performed the bioinformatic analyses, with help from GT, ZBa and NM. BK and DT drafted the article. MS and ZBo critically evaluated the paper. All named authors read and approved the final article.

