## Supplementary File 1 for "Combined Nanopore and Single-Molecule Real-Time Sequencing Survey of Human Betaherpesvirus 5 Transcriptome"

**This file contains Supplementary Figures S1-S4.**

**One other supplementary file is included as well:**

**Supplementary File 2.)** Excel table of the identified transcripts and their characteristics. The ORF compositions were assessed from the experimentally validated ORF list by Ster-Ginossar et al[1], considering either only the long ORFs or both the long and short ORFs, and also in combination with the *in-silico* predicted ORFs.

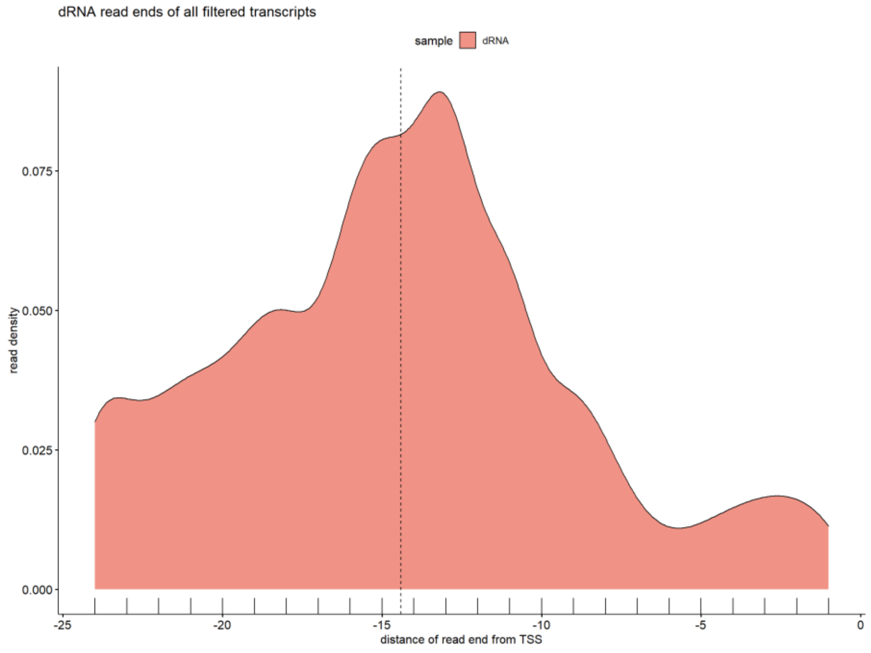


**Supplementary Figure S1.**) Density of the distances of the 5' ends of the reads from that of the transcript's (i.e., the zero position indicates the 5' end of the transcript) in the dRNA sample. Those reads are analyzed only, whose strand could be determined.

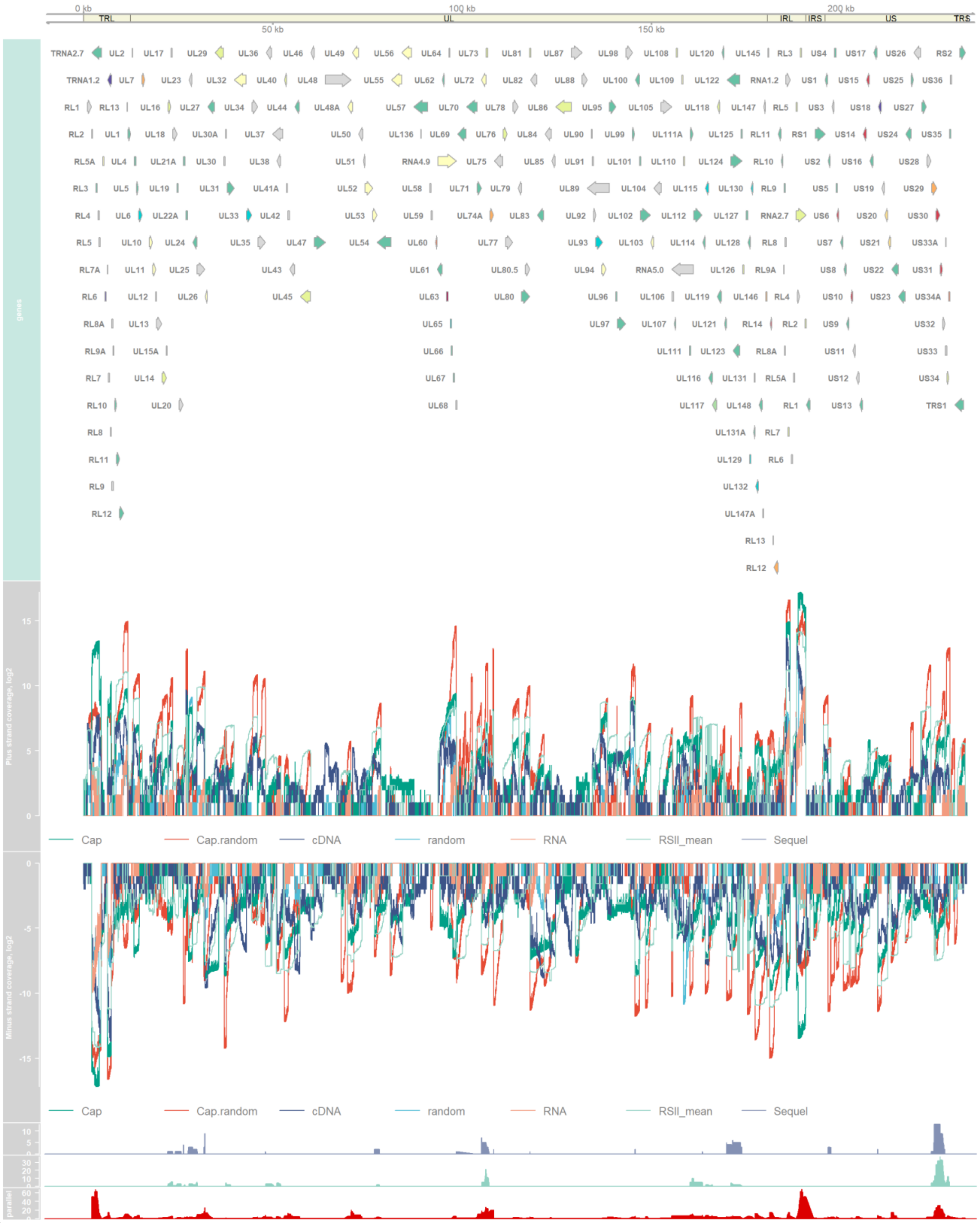


**Supplementary figure S2.**) HCMV genes (bottom panel), log2 coverages (middle panel) and transcriptional overlaps (calculated in a 10-nt window) of three different types (convergent, divergent, parallel) in the bottom panel. The genome is shown in two sections.

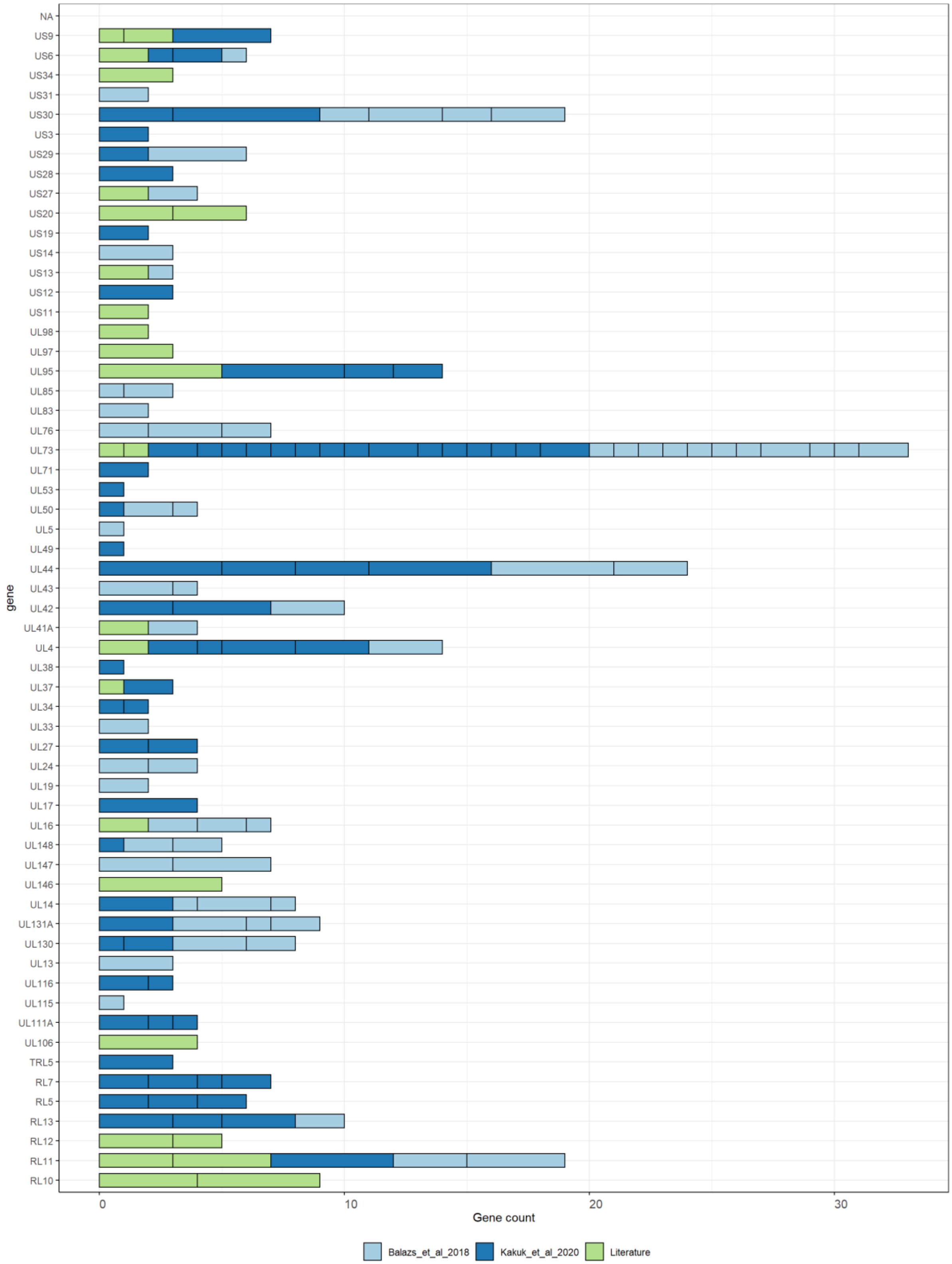


**Supplementary Figure S3.)** Genes that expressed polycistronic transcripts. Each bracket represents a polycistronic transcript, with its first gene shown on the Y-axis. The sizes of the brackets correspond to the number of genes that the respective transcript carries.

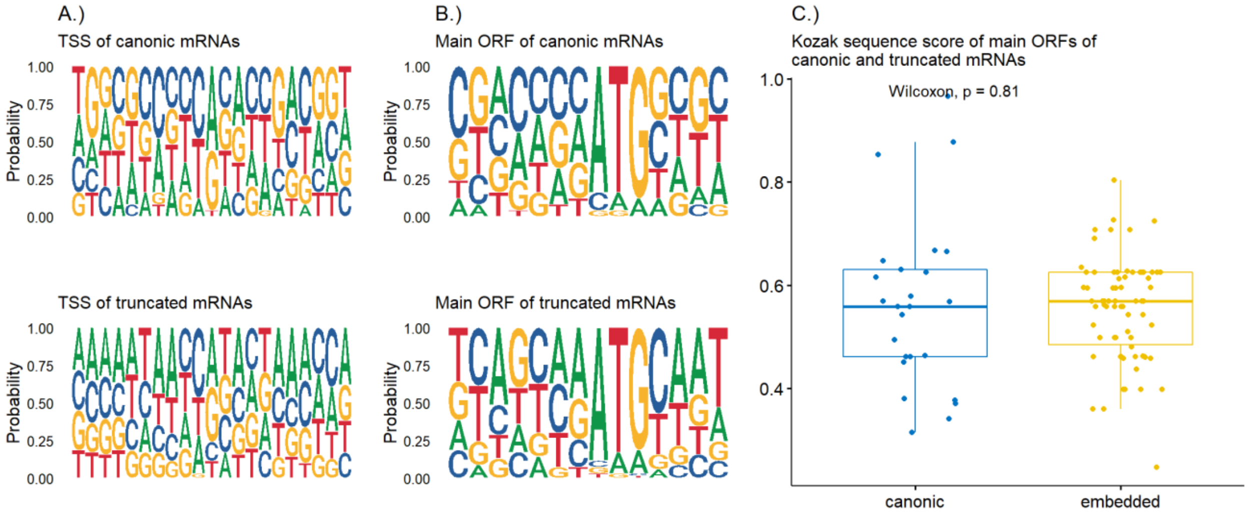


**Supplementary Figure S4. A.)** Weblogo of the sequences around the TSS of canonical and truncated transcripts (from -5 to +5 from the actual TES); **B.)** Weblogo of the Kozak consensus sequence the main ORFs of truncated and canonical; and **C.)** Kozak sequence score of the main ORFs of truncated and canonical transcripts.
